# A condensate dynamic instability orchestrates oocyte actomyosin cortex activation

**DOI:** 10.1101/2021.09.19.460784

**Authors:** Victoria Tianjing Yan, Arjun Narayanan, Frank Jülicher, Stephan W. Grill

**Affiliations:** Max Planck Institute of Molecular Cell Biology and Genetics, 01307 Dresden, Germany; BIOTEC, TU Dresden, 01307 Dresden, Germany; Max Planck Institute for the Physics of Complex Systems, 01187 Dresden, Germany; Center for Systems Biology Dresden, 01307 Dresden, Germany; Cluster of Excellence Physics of Life, TU Dresden, 01062 Dresden, Germany

**Author notes:** Correspondence: Correspondence and requests for materials should be addressed to A.N., F.J., S.W.G.

## Abstract

A key event at the onset of development is the activation of a contractile actomyosin cortex during the oocyte-to-embryo transition. We here report on the discovery that in *C. elegans* oocytes, actomyosin cortex activation is supported by the emergence of thousands of short-lived protein condensates rich in F-actin, N-WASP, and ARP2/3 that form an active micro-emulsion. A phase portrait analysis of the dynamics of individual cortical condensates reveals that condensates initially grow, and then switch to disassembly before dissolving completely. We find that in contrast to condensate growth via diffusion, the growth dynamics of cortical condensates are chemically driven. Remarkably, the associated chemical reactions obey mass action kinetics despite governing both composition and size. We suggest that the resultant condensate dynamic instability suppresses coarsening of the active micro-emulsion, ensures reaction kinetics that are independent of condensate size, and prevents runaway F-actin nucleation during the formation of the first cortical actin meshwork.

## Main text

Morphogenesis involves forces that are generated within the actomyosin cortical layer of cells^1, 2^. Improper cortical organization leads to an impairment of key cellular and developmental processes from as early as meiosis in oocytes to every subsequent cell division^3^. During meiotic maturation of oocytes, the actomyosin cortex transitions from inactive and non-contractile to active and tension-generating^4, 5^. This transition can generate a spectrum of actomyosin cortical structures and dynamics, including an actin cap in the mouse oocyte^6^, actin spikes in starfish oocytes^7, 8^, and waves of Rho activation and F-actin polymerization in *Xenopus*^9^. Organizing the first active actomyosin cortex requires the recruitment and assembly of various cortical components as well as the polymerization of actin filaments^10^. These processes have to be coordinated across the entire cell surface in order to generate a uniform actomyosin cortical layer. Here we ask how the formation of an active and tension-generating actomyosin cortex during meiotic maturation in oocytes is orchestrated.

The hermaphrodite nematode *Caenorhabditis elegans* (*C. elegans*) is a prime system for investigating actomyosin cortex formation during oocyte maturation^11–15^. In *C. elegans*, the onset of meiotic divisions and oocyte maturation coincides with ovulation and fertilization^11, 14, 15^. Oocytes are fertilized inside the hermaphrodite mother as they pass through the the sperm-containing organ – the spermatheca^11^. To understand how the formation of the first actomyosin cortex during oocyte maturation is orchestrated, we visualized F-actin in *C. elegans* oocytes containing Lifeact::mKate2. We observed that just before fertilization inside the mother, the oocyte cortical layer appears undeveloped with only sparse amounts of filamentous actin present (Figure 1 A left). In contrast, shortly after fertilization a highly dynamic and dense actomyosin cortical layer is present below the plasma membrane (Figure 1 A right, Movie S1). Importantly, we find that actomyosin cortex activation in the oocyte occurs through an intermediate stage that lasts approximately 10 minutes, results in a dynamic and contractile actomyosin cortical layer, and ends with the extrusion of the first polar body^16^ (Movie S1). Strikingly, this intermediate stage is characterized by the transient appearance of thousands of F-actin-rich condensates at the cortical layer (Figure 1 A). Here, we use the term condensate to refer to a dense assembly of specific molecular components maintained by collective molecular interactions. F-actin and its nucleators have previously been shown to form biomolecular condensates at the plasma membrane, and evidence for liquid-like properties was provided^17–21^. The F-actin rich condensates we observe are highly dynamic and inherently unstable. They appear stochastically and each disappears after approximately 10 seconds.

**Figure 1:**
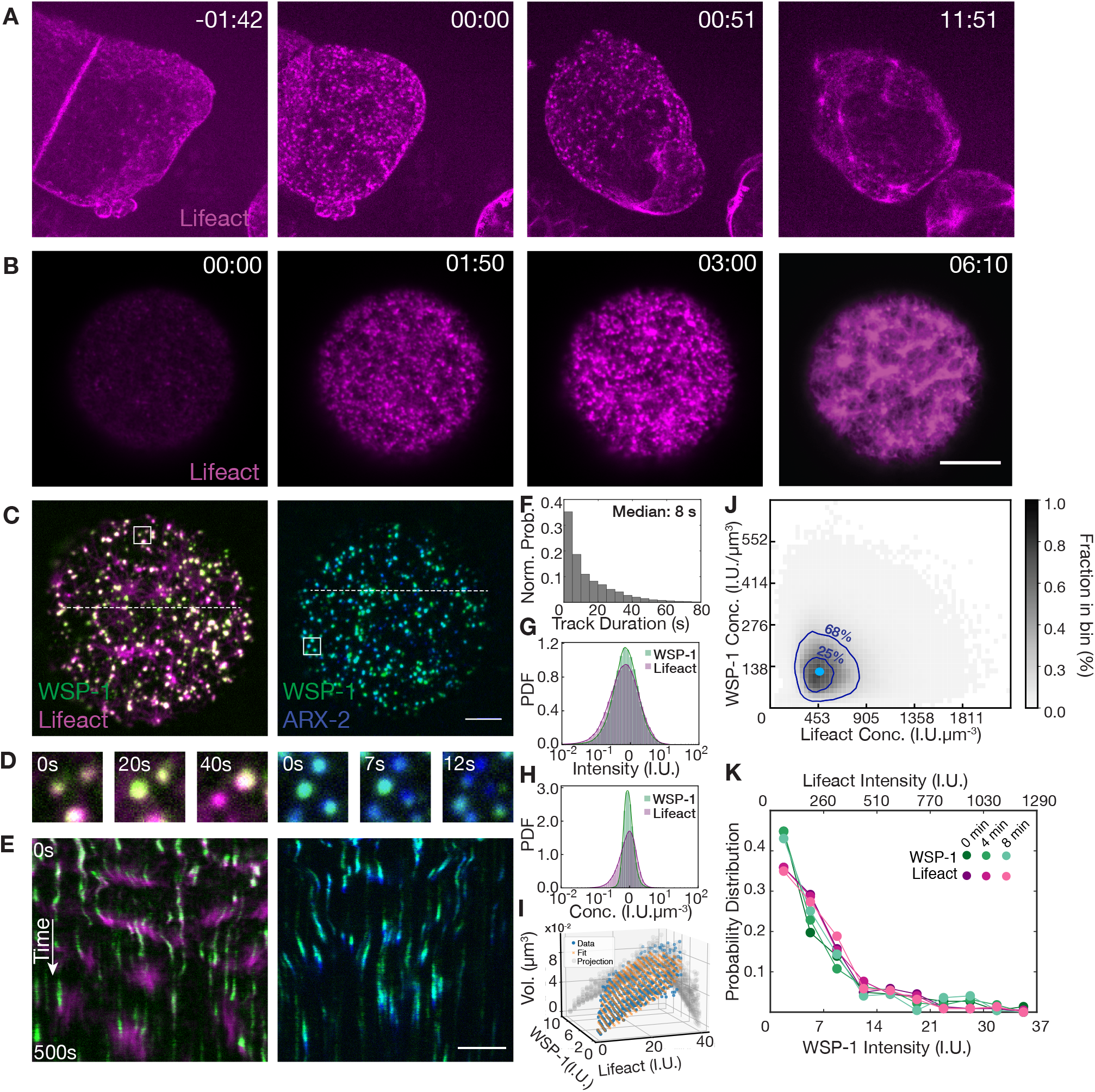
Actomyosin cortex formation at the oocyte to embryo transition proceeds through dynamic F-actin/WSP-1 cortical condensates. **A**, *In utero* microscopy images of the oocyte to embryonic transition in *C. elegans* at different times with respect to fertilization (min:sec). F-actin (Lifeact::mKate) in magenta, scale bars, 10 *μ*m (**A**-**E**). **B**, TIRF images of an isolated oocyte undergoing maturation. In both examples (**A, B**), a contractile cortex forms (rightmost image) following a stage characterized by the emergence of short-lived dynamic condensates rich in F-actin (two middle images). **C**, TIRF images of cortical condensates. Endogenous WSP-1::GFP in green (left), endogenous ARX-2::mCherry in blue (right). **D**, Compositional dynamics of condensates located within the respective white boxes in (**C**) over time, revealing that adjacent condensates can differ in their instantaneous dynamics. **E**, Condensate dynamics as revealed by kymographs obtained from the white dotted lines in (**C**). **F**, Normalized probability of condensate lifetime duration. **G**-**H**, Probability density distributions of Lifeact::mKate (magenta) and WSP-1::GFP (green) intensities (**G**) and concentrations (**H**) within condensates. **I**, Condensate volumes measured (blue) and calculated (orange) from the volume dependence on molecular content *υ*_*A*_*A* + *υ*_*W*_ *W*. **J**, Instantaneous concentrations of F-actin and WSP-1 within condensates from an ensemble of 36930 condensates from 9 oocytes. 68% and 25%of instantaneous condensate concentrations fall within the outer and inner dark blue contour line respectively. Light blue dot indicates the peak of preferentially maintained concentration pair of WSP-1 and F-actin in control oocytes. **K**, Normalized probability density distributions of WSP-1::GFP (green lines) and Lifeact::mKate (magenta lines) integrated condensate intensities are similar at 0, 4 and 8 minutes after the onset of oocyte maturation.

We next set out to investigate the nature of these transient F-actin rich condensates. To better observe their dynamics, we took advantage of the fact that oocytes isolated from the mother can mature in the absence of fertilization^22^. This allowed us to develop a TIRF assay for imaging cellular structures within ~ 200 nm of the cell membrane^23^ (Figure 1 B), to enable a quantitative study of actomyosin cortex formation in isolated oocytes at high spatial and temporal resolution (Movie S2). F-actin polymerization is organized by nucleation pathway members such as the branching nucleators N-WASP and ARP2/3 and the elongator Formin^24, 25^. We first investigated the presence of these components in cortical condensates. We used three strains labelling F-actin (by expressing Lifeact::mKate2) together with either endogenously labelled N-WASP (WSP-1::GFP), capping protein (CAP-1::GFP), or Formin (CYK-1::GFP)^26^. In addition, we used a strain that endogenously label both ARP2/3 (ARX-2::mCherry) and N-WASP (WSP-1::GFP). Besides F-actin, we identified N-WASP, ARP2/3 and the capping protein CAP-1^27^ (Figure 1 C, Figure S1, Movie S3 - 4) as components of cortical condensates, while the Formin CYK-1 was absent^16, 27^ (Figure S1). This demonstrates that cortical condensates contain molecules that mediate branched F-actin nucleation, similar e.g. to CD44 nano-clusters, dendritic synapses, and podosomes^28–30^. We also noted that during their ~ 10 second lifetime (Figure 1 F) cortical condensates were enriched first in N-WASP and ARP2/3, and only then F-actin accumulated before first losing N-WASP and ARP2/3 followed by F-actin (Figure 1 D,E). Given the time at which they appear and the fact that they contain molecules that mediate branched F-actin nucleation, we speculate that dynamic cortical condensates play a role in the formation of the first oocyte cortex.

We next asked if cortical condensates constitute a phase that coexists with the surrounding. Such a phase is characterized by material properties (such as density) that are intensive, ie. independent of volume. We used the strain that simultaneously labels F-actin and N-WASP to show that throughout their brief lifetime (Figure 1 E,F), cortical condensates varied over two orders of magnitude in both Lifeact (*A*) and WSP-1 (*W*) integrated fluorescence intensities (Figure 1 G). We estimated the volume of cortical condensates from the cross-sectional area determined by segmentation^31^ (see supplement), and found that for intensity stoichiometries *A/*(*A* + *W*) between ~ 0.65 and ~ 0.93, they occupied a volume *υ* well described by summing the volume contributions of F-actin *υ*_*A*_*A* and WSP-1 *υ*_*W*_ *W*, with volume coefficients *υ*_*A*_ = 1.54 10^−7^(±1 · 10^−8^) *μm*^3^*/IU* and *υ*_*W*_ = 2.34 10^−7^(±2 · 10^−8^) *μm*^3^*/IU* (where *IU* denotes total intensity units, see methods, Figure 1 I). This provides a relation between molecular content and volume, but does not imply that condensates are densely packed structures of only WSP-1 and F-actin. While cortical condensates varied over two orders of magnitude in integrated fluorescence intensities (Figure 1 G), the respective concentrations of WSP-1 and F-actin within the cortical condensates were significantly more restricted in their variation (Figure 1 H). This is also reflected in the emergence of a preferred pair of F-actin and WSP-1 concentrations maintained on average by the ensemble of cortical condensates (Figure 1 J). We conclude that on the one hand, cortical condensates are maintained far from equilibrium: they are highly dynamic and each disassemble after approximately 10s. On the other hand, cortical condensates display signatures of a multi-component condensed phase: they occupy a volume determined by their molecular content and exhibit a preferred pair of F-actin and WSP-1 concentrations distinct from their external environment^32, 33^. Hence, the ensemble of stochastically appearing, growing and subsequently dissolving cortical condensates effectively forms a chemically active micro-emulsion which, despite continuous turnover, maintains a steady size distribution that does not coarsen^34^ (Figure 1 K). Both the properties of a condensed phase and the mechanisms underlying its formation and dissolution can be revealed by a study of growth kinetics.

To study the growth kinetics of these cortical condensates, we quantified their compositions and volumes over time (Figure 2, see supplement). For a single representative cortical condensate, Figure 2 A,B shows the time evolutions of i) WSP-1 and F-actin total condensate intensity, ii) stoichiometry, and iii) volume (see supplement). For the example shown, WSP-1 precedes F-actin in both growth and loss, stoichiometry grows monotonically with time, and volume first increases and then decreases, and is well captured by summing volume contributions from F-actin and WSP-1. We noted that neighboring condensates followed similar trajectories in composition and volume despite forming stochastically and at different times (Figure 1 C-E). Thus, at a given time, neighboring cortical condensates which share their external environment can be at different stages of their internal life-cycle. We conclude that the growth kinetics of cortical condensates depend on their internal composition.

**Figure 2:**
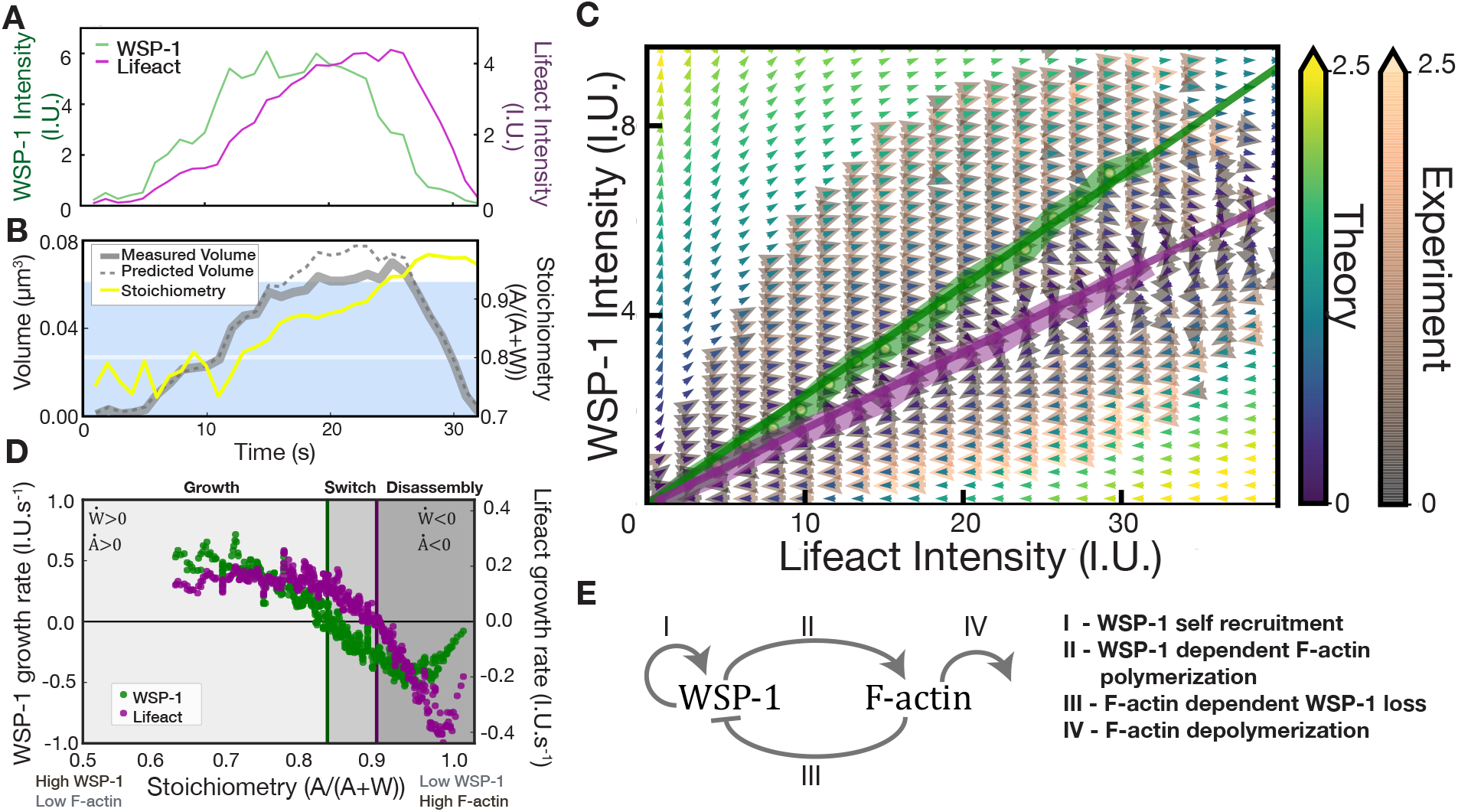
Mass flux phase portrait analysis of cortical condensate growth laws. **A**, Time trace of WSP-1::GFP (green line) and Lifeact::mKate (magenta line) total condensate intensities from a representative cortical condensates. **B**, Time trace of the volume measured (solid grey line), volume determined from the volume dependence on molecular content *υ*_*A*_*A* + *υ*_*W*_ *W* (dashed grey line), and stoichiometry 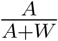 (yellow line) for the cortical condensate in (**A**). The region shaded in blue indicates the range of stoichiometry for which the *V* = *υ*_*A*_*A* + *υ*_*W*_ *W* accounts for measured volumes (see Figure S2). White line indicates estimated resolution limit. **C**, Mass flux phase portrait measured from 299165 time points of 36930 condensates from 9 oocytes (experiment, yellow-green-blue arrows), and calculated using the empirically determined growth laws (theory, orange-grey arrows). Color scales denote time rate change vector magnitudes. Thick lines indicate respective WSP-1 (green) and F-actin (magenta) nullclines from experiment, corresponding thin lines indicate nullclines from theory. **D**, Measured WSP-1 (green) and F-actin (magenta) growth rates as a function of stoichiometry display three distinct regimes separated by the WSP-1 nullcline at stoichiometry ~ 0.85, and the F-actin nullcline at stoichiometry ~ 0.9. **E**, Reaction motif underlying the structure of (**C**-**D**). The four processes are WSP-1 self-recruitment, WSP-1 dependent F-actin polymerization, F-actin dependent WSP-1 loss and F-actin depolymerization.

How does the internal composition of a cortical condensate influence its growth and shrink-age? To answer this question we developed a general method to quantitatively study compositional dynamics in an ensemble of multi-component condensates based on an analysis of the mass flux into the condensates (mass balance imaging^35^). For this, we quantified the time rate change of protein amounts within cortical condensates as a function of their internal F-actin and WSP-1 amounts. This time rate change of amounts is represented by a vector field, which defines average trajectories in the space of WSP-1 and F-actin amounts (Figure 2 C). Consistent with the representative example (Figure 2 A), average trajectories form loops that pass through three subsequent regimes: An early growth regime where condensates first grow in WSP-1 and subsequently in F-actin amounts, a switch regime where WSP-1 is lost while F-actin amounts still increase, and a disassembly regime with loss of both WSP-1 and F-actin. The nullcline of WSP-1 dynamics (green line in Figure 2 C), i.e. the WSP-1 amounts above which condensates grow and below which they shrink in WSP-1 content, reflects an F-actin dependent critical WSP-1 amount for WSP-1 growth. Stoichiometry is constant on lines that pass through the origin, hence the WSP-1 nullcline corresponds to a threshold stoichiometry of ~ 0.85. F-actin growth dynamics change from growth to shrinkage at a similar but slightly higher stoichiometry ~ 0.9 (the magenta line in Figure 2 C shows the F-actin nullcline). We conclude that cortical condensates become unstable and change from growth to disassembly in the switch regime between the two nullclines.

The three regimes (growth: above the WSP-1 nullcline, switch: between the two nullclines, disassembly: below the F-actin nullcline in Figure 2 C) are also visible when plotting WSP-1 and F-actin growth rates as a function of stoichiometry (Figure 2 D). Since the stereotypical compositional trajectories (Figure 2 A,B) involve a monotonic increase in stoichiometry with time, the xaxis of Figure 2 D also represents a progression through time. The dependence of growth rates on stoichiometry reveals the mutual regulation of WSP-1 and F-actin and can be depicted by the reaction motif shown in Figure 2E. Process I and II are mediated by WSP-1, process III and IV are mediated by F-actin. Process I corresponds to WSP-1 self-recruitment, evidenced by the fact that at low stoichiometry – corresponding to condensates consisting of mainly WSP-1 – the WSP-1 growth-rate is largest (Figure 2 D). Process II denotes WSP-1 dependent F-actin growth, reflected by a decrease of the F-actin growth rate as stoichiometry increases. This is most evident in the switch region of Figure 2 D. Process III denotes F-actin dependent loss of WSP-1, reflected by the fact that WSP-1 growth rates decrease with increasing stoichiometry. This suggests that F-actin counteracts the ability of WSP-1 to self-recruit, similar to previously reported negative feedback of F-actin on its nucleation via Rho^9^. Finally, process IV denotes F-actin depolymerization, reflected by the fact that F-actin is lost fastest at the highest stoichiometry (Figure 2 D). Further support for this reaction motif is provided by an analysis of WSP-1 and F-actin growth rates at constant WSP-1 and F-actin amounts (see Figure S3).

The shape of the measured phase portrait (Figure 2 C) and the shape of the growth rates 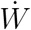 for WSP-1 and 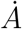 for F-actin as a function of stoichiometry (dots denote time derivatives; Figure 2 D) suggest the following empirical growth laws^36^ (see supplement, Figure S3, and Figure 2 C):

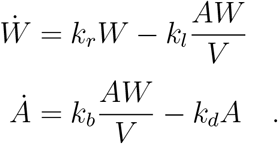

Here, WSP-1 self-recruitment depends linearly on *W* through the recruitment rate *k*_*r*_, consistent with the ability of WSP-1 to dimerize^37–39^ (process I). Interactions between F-actin and WSP-1 result in ARP2/3 mediated branched nucleation, and a subsequent increase in F-actin amounts^40^. This behaviour is captured by the term 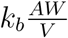, where *k*_*b*_ is a kinetic coefficient describing branching and condensate volume *υ* = *υ*_*A*_*A* + *υ*_*W*_ *W* depends on molecular amounts (see above, see supplement; process II). Branched nucleation coincides with a loss in WSP-1, this loss is captured by 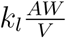 with the kinetic coefficient *k*_*l*_ describing the branching-dependent loss of WSP-1^41^ (process III). Finally, F-actin is lost with rate *k*_*d*_, consistent with severing and depolymerization^42^ (process IV). The mathematical form of all four terms are determined by the observation that the relative growth rates 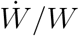 and 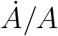 are linear functions of the effective F-actin volume fraction 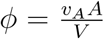 (see Figure 3E, F, see also discussion in supplement). Figure 3 E,F also allow us to estimate *k*_*r*_, *k*_*l*_, *k*_*b*_, *k*_*d*_. With these estimates, the the simple functions of the growth laws describe the experimental data remarkably well, and capture the entire mass flux phase portrait together with the composition dependent critical sizes as reflected by nullclines. (Figure 2 C).

**Figure 3:**
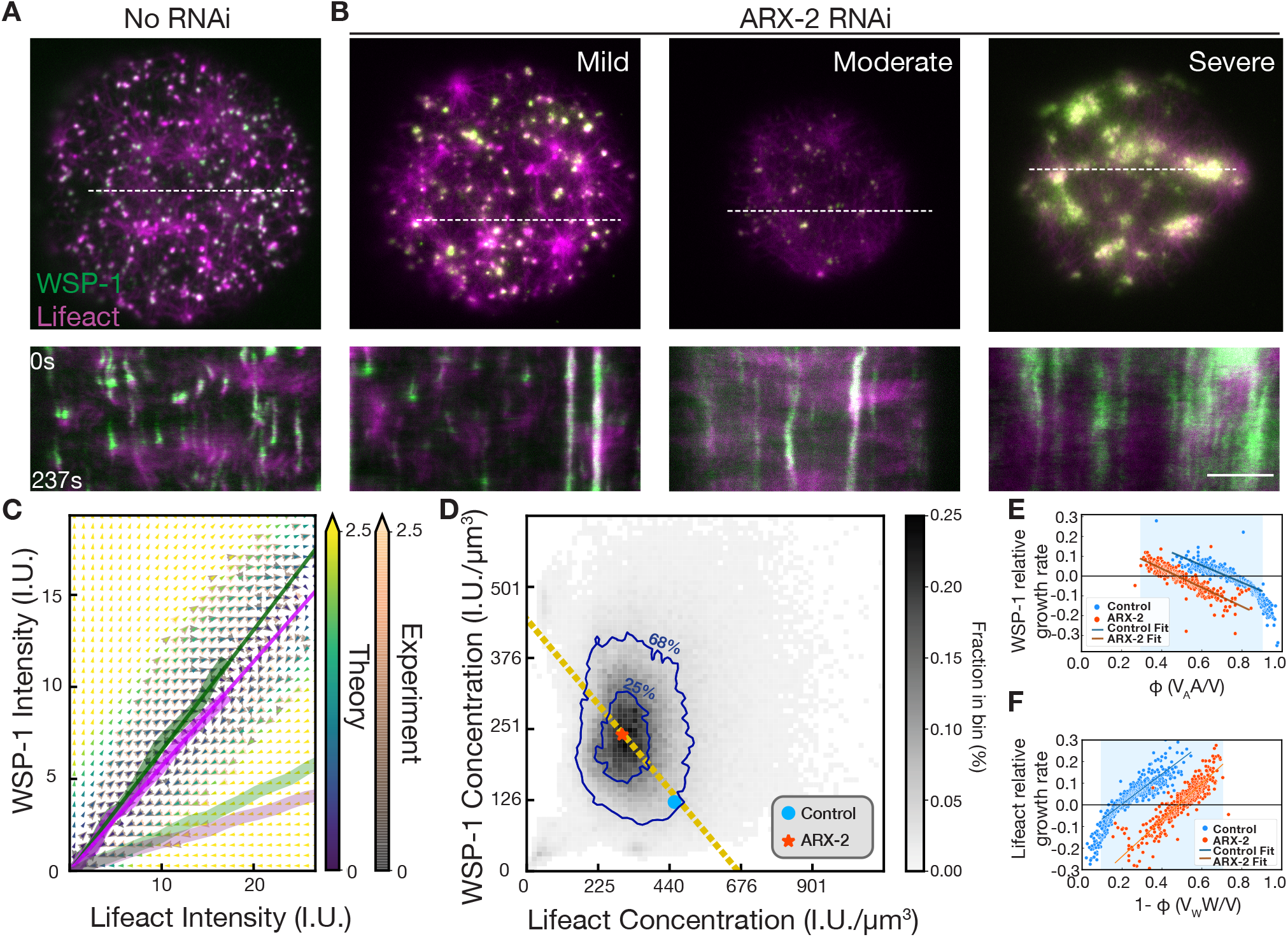
ARX-2 controls ensemble WSP-1/F-actin concentrations inside condensates by tuning condensate dynamics. **A**, Top, TIRF image of cortical condensates in an unperturbed control oocyte together with bottom, kymograph (determined along the dotted white line) to reveal temporal dynamics. **B**, Top, TIRF image of cortical condensates under mild (left, 16-17 hours RNAi feeding), moderate (centre, 17-18 hours RNAi feeding), and severe ARX-2 RNAi (right, *>* 18 hours RNAi feeding) together with bottom, respective kymographs (determined along the dotted white lines) to reveal temporal dynamics. Scale bars, 5 *μ*m. **C**, Experimental (yellow-green-blue arrows) and theory (orange-grey arrows) mass flux phase portrait of ARX-2 RNAi oocytes (7 oocytes, 13221 condensates and 180496 timepoints). Colors denote time rate change vector magnitudes. Thick lines indicate the corresponding WSP-1 (magenta) and F-actin (green) nullclines, shaded thick lines indicate nullclines from unperturbed control oocytes (Figure 2C). **D**, Histogram of instantaneous concentrations of F-actin and WSP-1 within condensates for the mild ARX-2 RNAi ensemble. 68% (25%) of instantaneous condensate concentrations fall within the respective blue contour lines. Blue dot and red star represent the preferentially maintained concentration pair for the ensemble of control and mild ARX-2 RNAi oocytes, respectively. Note that both lie on the orange dashed line of concentration pairs allowed by the volume dependence on molecular content. **E**-**F**, Linear dependence of relative WSP-1 (**E**) and F-actin (**F**) growth rates in the unperturbed control (blue) and mild ARX-2 RNAi case (red) on effective F-actin volume fraction *ϕ*(see supplement). Linearity holds within the region shaded in blue, indicating the range of stoichiometry for which the *V* = *υ*_*A*_*A* + *υ*_*W*_ *W* accounts for measured volumes (see Figure S2). Straight lines represent linear fits within the shaded region, yielding the parameters *k*_*r*_,*k*_*l*_ (**E**) and *k*_*b*_,*k*_*d*_ (**F**).

The WSP-1 nullcline *W*_*c*_(*A*) = *A*(*k*_*l*_ − *k*_*r*_*υ*_*a*_)*/k*_*r*_*υ*_*w*_ specifies a critical amount of WSP-1 above which WSP-1 amounts grow and below which WSP-1 amounts shrink. Notably, this critical amount plays a similar role as a critical droplet size for nucleation and growth, but here stems from biochemical reactions and not from condensation physics. The resulting growth laws exhibit a switch-like transition from growth to disassembly, representing a dynamic instability of condensates that is analogous to the dynamic instability of microtubules^43^.

To understand how the switch from condensate growth to condensate disassembly is orchestrated, we used RNAi to perturb the interplay between WSP-1 and F-actin. RNAi of actin regulators unrelated to branching such as RHO-1 (Rho GTPase), CYK-1, CDC-42, CHIN-1 (CDC-42 GAP), as well as multivalent adaptors VAB-1 (Ephrin receptor) and NCK-1 (Nck) did not affect cortical assembly dynamics^44–46^ (Figure S4). We focused on ARX-2 (ARP2 in the ARP2/3 complex in *C. elegans* ^47^), which mediates branched F-actin nucleation^10, 39^. Severe depletion of ARX-2 by RNAi (more than 20 hours of RNAi feeding at 20 °C) resulted in a loss of cortical condensates and dramatically altered cortical architecture (Figure 3 B right, Movie S5). On mildly depleting ARX-2 by RNAi (up to ~ 19 hours RNAi feeding at 20 °C; Figure 3 B left, Movie S6), oocytes showed reduced numbers of cortical condensates (average of 64±7 per oocyte as compared to 170±28 in unperturbed control oocytes, see Figure 3 A), which is consistent with a general reduction of F-actin branched nucleation ^48^.

An analysis of the remaining cortical condensates in ARX-2 depleted oocytes revealed that, in comparison to the unperturbed case compositional trajectories in mild ARX-2 RNAi are tilted towards the WSP-1 axis in the mass flux phase portrait (Figure 3 C). The mathematical form of the growth laws is maintained under mild ARX-2 RNAi, but the associated coefficients are changed (Figure 3 E,F). Mild ARX-2 RNAi reduced the rate of WSP-1 self-recruitment *k*_*r*_ by ~ 15%, and increased the coefficient *k*_*l*_ describing branching-dependent loss of WSP-1 by ~ 15%. In addition, the branching coefficient *k*_*b*_ remained essentially unchanged, while the F-actin loss rate *k*_*d*_ increased by a factor of ~ 2.75 (see supplement). A possibility consistent with mild ARX-2 RNA affecting WSP-1 recruitment but not branching-dependent F-actin generation is that WSP-1 inside condensates is bound to ARP2/3 (see supplement). The dominant effect of mild ARX-2 RNAi is the ~ 2.75-fold increase in the F-actin loss rate *k*_*d*_. This is consistent with previous findings that ARP2/3 protects F-actin from depolymerization *in vitro*^49^. In addition to the changes of coefficients, ARX-2 RNAi both reduced the average F-actin concentration and increased the average WSP-1 concentration in the condensate ensemble by factors of ~ 2 (Figure 3 D). Just like in the non-perturbed control, this pair of average concentrations falls on the line of constant total density of WSP-1 and F-actin combined (Figure 3 D yellow dashed line). We conclude that ARP2/3, largely through its impact on F-actin disassembly, governs the switch from condensate growth to condensate disassembly and determines the ensemble-averaged pair of internal concentration along the line of constant total density.

How do the growth kinetics lead to a specific pair of average internal concentrations and therefore a specific stoichiometry? To address this question we change variables from F-actin amount *A* and WSP-1 amount *W* to effective F-actin volume fraction *ϕ*= *υ*_*A*_*A/V* and condensate volume *υ*. Figure 4 A shows trajectories of condensates in the *ϕ*-*V* plane, revealing that the switch from condensate growth to shrinkage occurs at an effective F-actin volume fraction of ~ 0.8, corresponding to a stoichiometry of ~ 0.86. Notably, at this stoichiometry, the rate of change of condensate intensities, concentrations and volumes and thus also of stoichiometry is slowest (orange dotted lines in Figure 4 A,B,D), implying that the ensemble of dynamic condensates is governed by this slowly varying thus dominant stoichiometry. Hence, the peak of the concentration histogram (Figure 1 J, Figure 3 D) occurs at the point where the line of dominant stoichiometry intersects with the line of constant total density (Figure 4 D).

**Figure 4:**
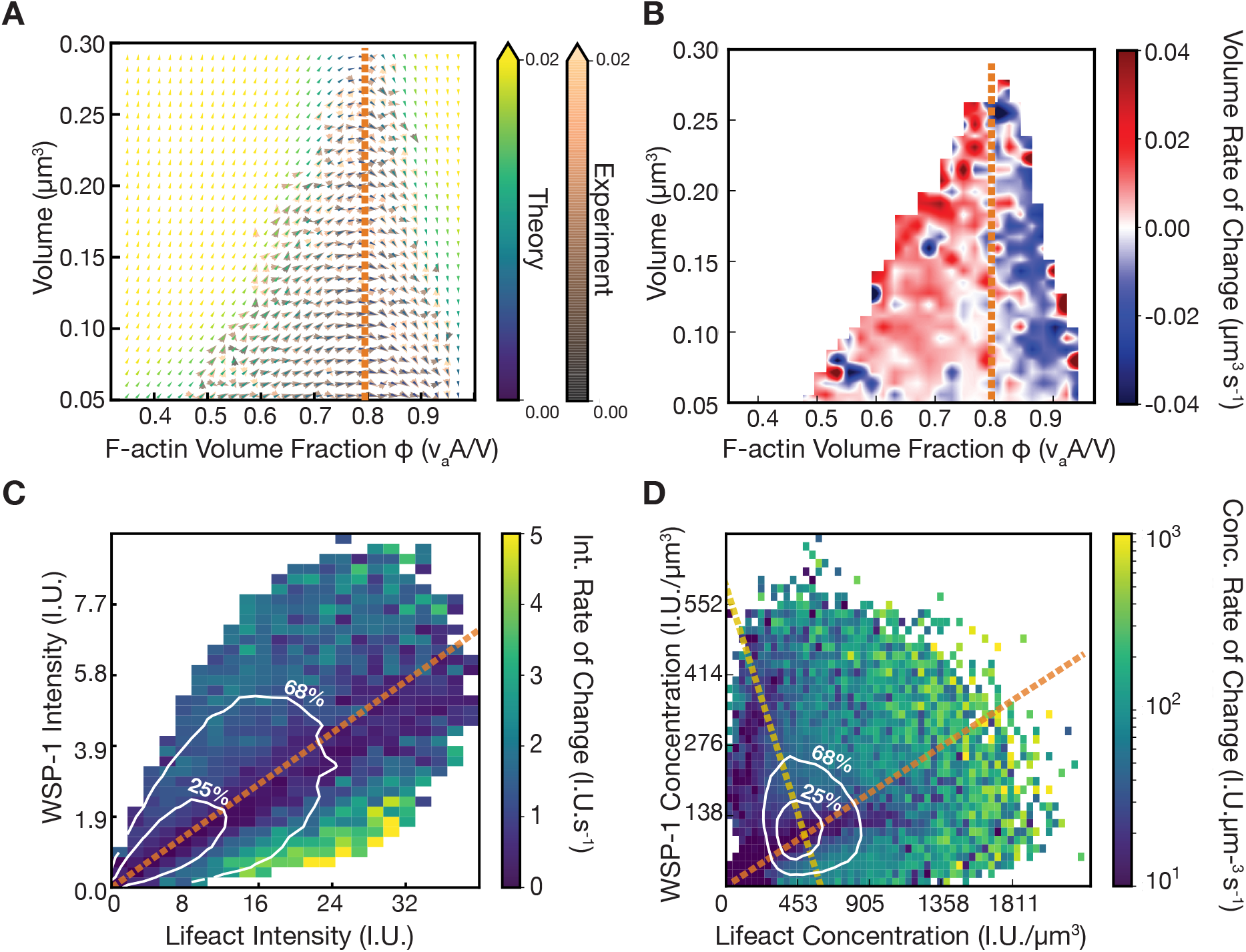
Volume-independent stoichiometry evolution sets internal concentrations. **A**, Volume-F-actin volume fraction phase portrait obtained by a change of variables of the data represented in Figure 2 C. Yellow-green-blue arrows, measured as in Figure 2 C. Orange-grey arrows, calculated using the empirically determined growth laws. Colors denote time rate change vector magnitudes. The thick line denotes the experimentally determined volume nullcline. The time evolution of effective F-actin volume fraction is independent of volume (see also supplement). **B**, Rate of change of volume as a function of instantaneous volume and effective F-actin volume fraction. Condensates switch from growth to shrinkage at an effective F-actin volume fraction of 0.8 (orange-dashed line), which corresponds to the region of slow kinetics within the switch region (Figure 2 D, 4 C-D). **C**, Mass flux phase portrait current magnitude. Contour lines (68%, 25%) depict the most commonly occupied total intensity values. The orange dashed line indicates the effective F-actin volume fraction corresponding to the center of the switch region (Figure 2 D), and coincides with lowest currents and slowest kinetics. **D**, Concentration flux phase portrait current magnitude. Contour lines (68%, 25%) depict the most commonly occupied concentration values and reflect the preferential maintenance of a pair of concentrations. This pair of concentrations lies at the intersect of the line of constant total density (yellow) and the line of dominant stoichiometry (orange) which corresponds to the orange lines in (**A**-**C**) and to the switch regions of Figure 2 C-D.

We also recognized that the time evolutions of the effective F-actin volume fraction 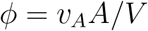 as well as WSP-1 and F-actin concentrations are independent of condensate volume (see supplement). Thus, condensate dynamics are intensive, which is consistent with mass action kinetics in well mixed systems. However, conventional mass action kinetics change reactant concentrations at constant volume, but usually do not involve assembly and disassembly as is the case here. Note that intensive condensate dynamics are not consistent with conventional kinetics of nucleation and growth of liquid-like condensates, where assembly rates depend on condensate size^50, 51^. This reveals that cortical condensates exhibit an unconventional chemical kinetics where mass action governs assembly and disassembly, and therefore the effect of mass-action dynamics on concentrations in condensates is modified (see supplement). Note, however, that even though cortical condensates do not assemble via classical nucleation and growth, the intensive condensate dynamics reveal that the condensate material behaves as a well-mixed phase with size-independent properties. Intensive reaction dynamics are expected to arise in situations where the time for diffusion across the condensate is shorter than the typical time associated with a chemical reaction. The condensate dynamic instability limits cortical condensate size. Therefore, reaction dynamics remain intensive and the resultant chemically active micro-emulsion maintains a steady-state size distribution (Figure 1 K)^34^.

To conclude, cortical condensates represent a new type of non-equilibrium biomolecular condensate that assembles and disassembles via a dynamic condensate instability governed by mass action chemical kinetics. They recruit molecules that drive branched nucleation of F-actin, and support the activation of the actomyosin cortex. The dynamic instability of cortical condensates is similar to the dynamic instability of growing and shrinking microtubules^52, 53^, but arises in a bulk assembly that forms a phase. We suggest that the formation and subsequent dissolution of cortical condensates via a condensate dynamic instability serves to control autocatalytic F-actin nucleation and prevents runaway growth during the activation of the first cortical actin meshwork in the *C. elegans* oocyte.

## Methods

See supplemental materials.

## Acknowledgements

S.W.G. was supported by the DFG (SPP 1782, GR 3271/2, GR 3271/3 and GR 3271/4) and the European Research Council (grants no. 742712, H2020-MSCA-ITN-2015). V.Y. acknowledges Marie Skłodowska-Curie Actions (grant no. H2020-MSCA-ITN-2015) for funding and support. A.N. thanks the ELBE program of MPI-PKS and MPI-CBG for funding and support. We are grateful to 2017 and 2019 MBL Physiology course faculty and students, in particular A. Chakrabarti, M. Dietrich, G. Martinez, M. Mirvis, and Q. Yu. We thank A. Bhatnagar, S. Choubey, K. Crell, E. Garner, J. Geisler, N. Goehring, M. Holmes, A. Honigman, T. Middlekoop, A. Mukherjee, J. Reich, G.Squyres, and T. Wiegand for experimental help and discussions. We are grateful to J. Brugués, O. Campa’s, P. Goünczy, and W. Grill for critical comments on the manuscript.

