## Supplemental Material Methods and Notes for "A condensate dynamic instability orchestrates oocyte actomyosin cortex activation"

### 1 Materials and Methods

***C. elegans* maintenance and strains** The *C. elegans* strains used in this study were cultured on nematode growth media plates, seeded with OP50 *E. coli*. Strains were incubated in 20 °C until imaging at 22-24 °C. The following transgenic strains were used:

| Strain | Description | Source |
|---|---|---|
| SWG022 | Lifeact::mKate2 PH-domain::GFP | Grill lab |
| SWG007 | Lifeact :: mKate2/nmy-2 :: GFP | Grill lab |
| MG589 | GFP::utrophin | Glotzer lab |
| SWG251 | GFP::utrophin;Lifeact :: mKate2 | This study |
| GOU1861 | GFP::wsp-1a; arx-2::TagRFP | Gift from the Ou lab |
| SWG61 | CAP-1::eGFP CRISPR; Lifeact::mkate2 | Grill lab |
| SWG62 | CYK-1:eGFP CRISPR; Lifeact::mkate2 | Grill lab |
| SWG197 | ARX-2::gfp V;Lifeact::mKate2; | This study |
| SWG198 | GFP::wsp-1 IV;Lifeact::mKate2 | This study |

GFP::Utrophin and PH-domain::gfp were generated by knock-in transgenesis, Lifeact::mKate is generated with Mos1-mediated Single-Copy Insertion (MosSCI), all other fluorescent labelling of proteins are expressed from the endogenous locus with CRISPR mutagenesis, from both the work of Anne Cecile Reymann, and generous gift from Guangxu Ou lab.

***In utero* confocal imaging** Adult *C. elegans* hermaphrodites were anaesthetized for 3 minutes with 0.5% Tetramisole (Sigma-Aldrich T1512) in M9 buffer, then placed on an 18x18 mm high-precision coverslip precoated with 0.1% Poly-L-Lysine (Sigma-Aldrich P8920). Images were obtained on a spinning disk confocal microscope (Zeiss C-Apochromat, 63x/1.2 NA objective, with Hamamatsu Orca-Flash 4.0 camera). Images were acquired at 3 second intervals with 3 Z-slices (totalling 1.5 $\mu$ m). The excitation lasers used were 488 and 568 nm, and the exposure time was 100ms for each channel.

**Intensity units** Intensity units (I.U.) for both the F-actin and WSP-1 channels correspond to the camera readings divided by 10,000 for ease of presentation and as a proxy for molecular amounts.

**Oocyte TIRF imaging** Oocytes of adult hermaphrodites were dissected as described by reference [1], and imaged in Shelton's growth media[2]. 1% 20 $\mu$m polystyrene beads were used in the growth media to act as a spacer for imaging. Cells imaged in TIRF microscopy were placed on ethanol cleaned 18x18 mm high-precision coverslips. TIRF microscopy of oocytes was conducted on a Nikon Ti microscope equipped with a TIRF turret. The alignment and collimation of TIRF lasers were performed at the beginning of each imaging session. Movies were acquired at 1s intervals, with 100ms exposure time for each channel. The laser lines used were 488nm, and 568nm lasers, for GFP, mKate/mCherry, and JF647 respectively. The images were acquired on an Andor iXon-Ultra EMCCD with 1.5x optical zoom. Nikon Perfect Focus was used to maintain focus of the

TIRF plane, throughout the duration of imaging. A 488 nm laser was used with a 525/50 bandpass emission filter and a 561 nm laser was used with a 568 nm long pass filter, for imaging GFP and mKate respectively.

**Kymographs** Kymographs were used to illustrate dynamics in time. Line kymographs were generated in Fiji/ImageJ.

**Oocyte chemical inhibitor treatments** Chemical inhibitors were directly added to Shelton's growth media during the dissection stage, as they are cell-permeable, and the oocytes have no eggshell. CK-666 was used at 10 $\mu$M.

**RNAi Feeding** Knockdowns of proteins were achieved by RNAi clone feeding to L4 stage *C. elegans*. RNAi clones were acquired from the Ahringer lab RNAi library, and seeded onto a feeding plate (NGM agar containing 0.5 mM to 2 mM IPTG and 25$\mu$g/ml carbenicillin). L4 stage worms were placed onto feeding plates and incubated at 20°C for 15 to 30 hours, depending on the intended strength of depletion.

**Phase portrait analysis pipeline** Microscopy images used for analysis were first exported using Fiji/ImageJ. Background subtraction of the camera noise is performed with a custom Matlab script. Intensities from both GFP and mKate channels were summed prior to segmentation, in order to capture full duration of tracks. Puncta were segmented with Ilastik[3]. Intensities of segmented puncta were measured using Matlab regionprops. Tracking of the puncta traces were performed with Fiji Trackmate (Simple LAP tracker)[4]. Track intensity parsing, as well as phase portrait generation from data, and nullcline calculation were performed with Matlab and Python custom written code. Volumes shown in Figure 4 are above the estimated lateral resolution of $\sim 200nm$.

**Parameter estimation** Least squares fitting of the linear dependence of relative growth rate on stoichiometry ($\frac{\dot{W}}{W} = k_r - k_l \frac{A}{A+W}$ and similarly for $A$) (Figure 3 E,F) was implemented in *Python (Lmfit)*. Measured volume coefficients ($V = v_A A + v_A W$) were determined from the measured spatial data for clusters larger than the estimated lateral resolution $\sim$ 200 nm and the measured F-actin and WSP-1 intensities by the 'leastsq' method of Python Lmfit. These results were used for the generation of Figure 2 C, Figure 3 C, Figure 4 A and Figure S5A. They were:

$k_d = 0.15(\pm 0.01)$ ($k_d = 0.39(\pm 0.01)$ with ARX-2 RNAi) and $k_r = 0.54(\pm 0.01)$ ($k_r = 0.45(\pm 0.01)$ with ARX-2 RNAi), $k_l/v_A = 0.73(\pm 0.03)$ ($0.94(\pm 0.04)$ with ARX-2 RNAi) and $k_b/v_W = 0.78(\pm 0.02)$ ($0.81(\pm 0.02)$ with ARX-2 RNAi). Volume $V = v_A A + v_A W$ with $v_A = 1.54 \cdot 10^{-7}(\pm 1 \cdot 10^{-8})$ ($2.3 \cdot 10^{-7}(\pm 2 \cdot 10^{-8})$ with ARX-2 RNAi) and $v_W = 2.34 \cdot 10^{-7}(\pm 2 \cdot 10^{-8})$ ( $3.7 \cdot 10^{-7}(\pm 2 \cdot 10^{-8})$ with ARX-2 RNAi)

### 2 Dependence of condensate volume on composition

In the main text we showed that the volume of a cortical condensate is determined by a linear combination of its F-actin and WSP-1 content as reflected in the integrated fluorescence intensities (Figure 1 I). Approximately $80\%$ of events lie in F-actin volume fraction range from 0.6 to 0.9 (corresponding to intensity based stoichiometry $\frac{A}{A+W}$ from $\sim$ 0.65 to $\sim$ 0.93). For all events in this range, the measured volume follows $V = v_A A + v_W W$. All stoichiometries discussed in Figures 2-4 lie in this range. For the minority of assemblies beyond this range, the deviation between the growth laws and the data is of the same order as the deviation in volumes (Figure S2) - which provides further evidence for mass action determined growth rates.

### 3 Condensate stoichiometry and effective F-actin volume fraction

In the main text, based on measured intensities alone, we define stoichiometry S= $\frac{A}{A+W}$. This also corresponds to a compositional fraction of F-actin. Based on both measured intensities and the measured volumes, we define the effective F-actin volume fraction $\phi = \frac{v_A A}{v_A A + v_W W}$. Given our estimation of the volume contribution coefficients $v_A$ and $v_W$ (see Section 2 and Figure 1 I), the two definitions are related by $\phi = \frac{S}{S - \frac{v_W}{v_A}(1-S)}$. Thus the threshold intensity stoichiometry of $\sim .86$ (between the nullclines at $S = 0.85$ and $S = 0.89$ in Figure 2) corresponds to F-actin volume fraction of $\phi = 0.8$ (as seen in the orange line in Figure 4).

### 4 Determination of empirical condensate growth laws and their interpretation

The experimental data enforces constraints on the dependencies of F-actin and WSP-1 growth rate on internal composition ($A$ and $W$ amounts). These constraints are simultaneously captured in the linear dependence of the measured relative growth rates ($\frac{\dot{A}}{A}$, $\frac{\dot{W}}{W}$) as a function of the effective F-actin volume fraction ($\phi = v_A A/V$) (main text Figure 3 E,F). The positive intercept in the case of $\frac{\dot{W}}{W}$ as a function of $\phi$ (Figure 3 E) fixes both the existence and form of the WSP-1 self recruitment term ($+k_r W$) while the negative intercept in the case of $\frac{\dot{A}}{A}$ as a function of $1 - \phi$ (Figure 3 F) fixes the existence and form of the F-actin loss term ($-k_d A$). Additionally, the fact that the relative growth rates for F-actin and WSP-1 change linearly with F-actin volume fraction $\phi$ (or $1-\phi$) fixes the existence and form of the cross terms ($+k_b\frac{AW}{V}$ and $-k_l\frac{AW}{V}$). Thus, the experimentally observed linearity of the functions $\dot{W}/W = k_r - \frac{k_f}{v_A}\phi$ and $\dot{A}/A = \frac{k_l}{v_W}(1-\phi) - k_d$ with respect to $\phi$ completely determines the form of all four terms in the growth laws.

This mathematical form is also reflected in the phase portrait slices of Figure S3. The dependencies of $\dot{W}$ and $\dot{A}$ on $A$ and $W$ include contributions from two terms each : $k_rW$ and $-k_l\frac{AW}{V}$ in the case of $\dot{W}(A,W)$ and $-k_dA$ and $+k_b\frac{AW}{V}$ in the case of $\dot{A}(A,W)$. Figure S3 D shows that at low WSP-1 levels, F-actin is lost with a rate that grows linearly with $A$ ( $-k_dA$ ). At the low WSP-1 levels of Figure S3 D the contribution from the $+k_b\frac{AW}{V}$ term is minimal allowing the linear dependence to be directly visible. Similarly the form of the cross dependencies ($-k_l\frac{AW}{V}$ and $+k_b\frac{AW}{V}$ ) are reflected in the linear-to-saturating behavior of the Figure S3 B,C just as the functions $-k_l\frac{AW}{V}$ and $+k_b\frac{AW}{V}$ should behave with $V = v_AA + v_WW$. Finally the non monotonic behaviour of Figure S3 A reflects the competing contributions of both WSP-1 self recruitment and F-actin dependent WSP-1 loss. Importantly, these dependencies recapitulate the full phase portraits in both control (Figure 2 C, Figure 4 A) and ARX-2 RNAi (Figure 3 C, Figure S5) conditions. Note that the growth laws in both control and ARX-2 RNAi conditions simultaneously recapitulate phase portraits resulting from two distinct measurements in two distinct coordinate systems - intensity measurements for the mass flux phase portrait of Figure 2 C (3C for RNAi) and volume measurements for the volume-volume fraction phase portrait of Figure 4 A (S5 for RNAi).

### 5 Condensate growth laws under perturbation of ARX-2 RNAi

The changes of rate coefficients on ARX-2 RNAi suggest biochemical interpretations. First The F-actin independent WSP-1 self-recruitment rate $k_r W$ reduces by approximately $15\%$ [Figure 3 E] on ARX-2 RNAi suggesting that ARX-2 is involved in WSP-1 self-recruitment and potentially that ARX-2 bound WSP-1 is preferentially recruited to the condensates. Secondly, if ARX-2 bound WSP-1 are enriched in the condensates, depletion of ARX-2 should affect the recruitment of WSP-1, and the branching rate of recruited ARP2/3 are not expected to change. Consistent with this, the $(+k_b \frac{AW}{V})$ term is unchanged on ARX-2 RNAi. Thirdly, WSP-1 is known to detach from ARP2/3 on branching[5]. As the rate of branching reactions ($k_b$) is essentially unchanged, the $\sim 15\%$ increase in WSP-1 loss rate ($k_l$) suggests that the escape time of WSP-1 molecules post-branching is partially modulated by free ARP2/3 within the cluster. For instance, the modulation potentially proceeds by rebinding of WSP-1 with ARX-2 and additional branching cycles. Finally as discussed in the main text, ARX-2 RNAi led to a $\sim 3$ fold increase in the depolymerization rate $(-k_d A)$. This is consistent with a a well-established propensity of ARP-2/3 to bind and prevent depolymerization at the pointed ends of F-actin filaments[6]. These interpretations are schematically depicted in Figure S6.

### 6 Volume independent condensate dynamics

Intensive properties defining a phase are those properties that are independent of volume (eg. concentration). By intensive condensate dynamics we mean the volume-independent time-evolution of the intensive properties defining the condensate phase. In the main text we stated that the growth laws imply a volume independent time-evolution of F-actin volume fraction $\phi = v_A A/V$, WSP-1 concentration $W_c = W/V$ and F-actin concentration $A_c = A/V$. Below we show that the time evolution of $W_c, A_c$ and $\phi$ depends only on internal concentrations and not on condensate size.

We start from the growth laws as extracted from the experimental data

$$\dot{W} = k_r W - k_l \frac{AW}{V} \tag{1}$$

$$\dot{A} = k_b \frac{AW}{V} - k_d A \quad , \tag{2}$$

with $V = v_A A + v_W W$, and k's as defined in main text. We first divide the equations by volume V to get

$$\frac{\dot{W}}{V} = k_r W_c - k_l A_c W_c \tag{3}$$

$$\frac{\dot{A}}{V} = k_b A_c W_c - k_d A_c \quad , \tag{4}$$

Where $A_c = A/V$ and $W_c = W/V$ are the internal concentrations. As condensate volume is not constant, $\dot{W}_c = \dot{W}/V - W_c \dot{V}/V$ and similarly $\dot{A}_c = \dot{A}/V - A_c \dot{V}/V$. Substituting in the growth laws we get :

$$\dot{W}_c = k_r W_c - k_l A_c W_c - W_c \frac{\dot{V}}{V} \tag{5}$$

$$\dot{A}_c = k_b A_c W_c - k_d A_c - A_c \frac{\dot{V}}{V} \quad , \tag{6}$$

From the definition $V = v_A A + v_W W$ and the growth laws for $\dot{A}, \dot{W}$ it follows that

$$\frac{\dot{V}}{V} = v_A (k_b A_c W_c - k_d A_c) + v_W (k_r W_c - k_l A_c W_c) \quad , \tag{7}$$

is a function of only concentrations. Thus, the time evolutions of $W_c, A_c$ on the left hand side, are purely determined by concentrations on the right hand side and are independent of condensate volume i.e. intensive.

For the case of effective F-actin volume fraction $\phi = v_A A/V$ we first note that $1 - \phi = v_W W/V$ since $V = v_A A + v_W W$. This allows us to express $\dot{W}$ and $\dot{A}$ as functions of $V, \phi$ :

$$\dot{W} = \frac{k_r}{v_W} V(1-\phi) - \frac{k_l}{v_A v_W} V\phi(1-\phi) \tag{8}$$

$$\dot{A} = \frac{k_b}{v_A v_W} V\phi(1-\phi) - \frac{k_d}{v_A} V\phi \quad . \tag{9}$$

The time rate change of $V$ and $\phi$ follow from their definitions above. We get:

$$\dot{V} = v_A \dot{A} + v_W \dot{W} \tag{10}$$

$$\dot{\phi} = \frac{v_A}{V}\dot{A} - \frac{v_A A}{V^2}(\dot{A}\partial_A V + \dot{W}\partial_W V) \quad . \tag{11}$$

Substituting for $\dot{A}$ and $\dot{W}$ in terms of V and $\phi$ gives us the time rate change of $V$ and $\phi$ as functions of volume and stoichiometry:

$$\dot{V} = V\left[k_r(1-\phi) - k_d\phi + (\frac{k_b}{v_W} - \frac{k_l}{v_A})\phi(1-\phi)\right] \tag{12}$$

$$\dot{\phi} = (\frac{k_b}{v_W} - k_d - k_r)\phi + (k_d + k_r - \frac{2k_b}{v_W} + \frac{k_l}{v_A})\phi^2 + (\frac{k_b}{v_W} - \frac{k_l}{v_A})\phi^3 \quad . \tag{13}$$

Thus the evolution of F-actin volume fraction is given by a polynomial in $\phi$ and is independent of volume.

### 7 Mass action kinetics during assembly and disassembly of cortical condensates

The law of mass action is typically applied to chemical reactions in a well-mixed container of fixed volume and states that the rate of a chemical reaction is proportional to the product of the activities or concentrations of the reactants (see for instance reference [7]). For example, consider $n_A$ molecules of type A and $n_B$ molecules of type B, whose one-step reaction produces molecule C, within a container of constant volume V. In this case, changes in the number of product molecules $n_c$ arise from the rate at which molecules of type A and molecules of type B meet and react. Since the solution is well-mixed, the number of type A molecules within reaction-range of type B molecules at any instant is just the probability that the random placement of $n_A$ molecules of type A and $n_B$ molecules of type B within a Volume V results in such an overlap. This probability is proportional to $n_A n_B / V$ and thus,

$$\dot{n_C} = k \frac{n_A n_B}{V} \quad , \tag{14}$$

where k is the kinetic coefficient for production of C from proximity of A to B. Since the volume of the reaction container is a constant, on dividing by this volume, we get the familiar expression for the law of mass action for concentration changes

$$\dot{C_c} = k A_c B_c \quad , \tag{15}$$

where $C_c, A_c$ and $B_c$ denote the concentrations $n_C/V$, $n_A/V$ and $n_B/V$ respectively and the resulting time-evolution of $C_c$ is intensive - i.e. depending only on the intensive quantities $A_c, B_c$ and independent of volume V.

In the case of cortical condensates, the volume is not constant as the reaction container itself evolves with molecular amounts. A key result of our work is that mass action kinetics describe cortical condensate evolution at any instantaneous volume. To see this, first consider the branching reaction with two reactants $A$ and $W$ at concentrations $A_c = A/V$ and $W_c = W/V$. The rate at which branching reactions lead to a loss of WSP-1 ($\dot{W} = -k_l AW/V$) or addition of F-actin ($\dot{A} = k_b AW/V$) are proportional to collision rates of the two reactants $A$ and $W$ in a well mixed container as they have the form of the product of concentrations $A_c W_c = AW/V^2$ summed over the instantaneous condensate volume V. Second, consider WSP-1 self-recruitment rate ($k_r W$), WSP-1 self recruitment can be considered a reaction between internal WSP-1 at the concentration $W_c$ and a constant permeating bath of recruitable WSP-1. Then, this term has the form of the product of internal WSP-1 concentration with the constant bath concentration and summed over the instantaneous condensate volume V. Lastly, the F-actin loss rate ($-k_d A$) requires a single reactant ($A$) to depolymerize and is also proportional to concentrations of this reactant $A_c$ summed over the instantaneous condensate volume.

Thus the growth laws:

$$\dot{W} = k_r W - k_l \frac{AW}{V} \tag{16}$$

$$\dot{A} = k_b \frac{AW}{V} - k_d A \quad , \tag{17}$$

are consistent with mass action kinetics in the instantaneous condensate volume V. Dividing by V, we get

$$\dot{W}/V = k_r W_c - k_l A_c W_c \tag{18}$$

$$\dot{A}/V = k_b A_c W_c - k_d A_c \quad . \tag{19}$$

However, as V varies with time, the left hand sides are not simply the rate of change of WSP-1 and F-actin concentrations $\dot{W}_c$, $\dot{A}_c$. Instead, since $\dot{W}_c = W/V$ and $\dot{A}_c = A/V$ we get

$$\dot{W}_c = k_r W_c - k_l A_c W_c - W_c \dot{V}/V \tag{20}$$

$$\dot{A}_c = k_b A_c W_c - k_d A_c - A_c \dot{V}/V \quad . \tag{21}$$

The additional terms $W_c \dot{V}/V$ and $A_c \dot{V}/V$ on the right hand side arise from volume changes. In order for the evolution of internal concentrations to be volume independent (intensive), $\dot{V}/V$ must be volume independent.

In the case of cortical condensates, the number of molecules evolve according to mass action rates and determine the volume, the term $\dot{V}/V$ is a function of only internal concentrations (see Equation 7 above). The time evolution of $A_c$ and $W_c$ within cortical condensates is therefore intensive as shown in the previous section.

Finally, in the main text we state that 'intensive condensate dynamics are not consistent with conventional kinetics of nucleation and growth of liquid-like condensates'. As an example of this, note that if WSP-1 were accumulating via diffusion limited nucleation and droplet growth from its supersaturated vapor, the WSP-1 self recruitment term would have the size-dependent form $\dot{W} \sim aW^{1/3} - b$ (equation 2.53 in reference[8]) with the constants $a$ and $b$ depending on thermodynamic parameters such as external diffusion constant, supersaturation and a capillary length. In this case, the cross terms $k_l \frac{AW}{V}$ and $k_b \frac{AW}{V}$ and depolymerization term would remain of mass action form and yet the time-evolution of $A_c$, $W_c$ and $\phi$ would not be intensive.

### 8 Requirements for achieving volume-independent condensate dynamics

We may generalize the discussion in section 6 to clarify the requirements for achieving intensive condensate dynamics. We consider an arbitrary scheme of chemical reactions involving $n$ reactants generating a product in a container of varying volume $V$. The system is *well-mixed* if the system is effectively homogenized by diffusion on the shortest reaction time scale. This requires that the timescale for diffusion across the container is faster than any reaction. This requirement is satisfied when the container size is smaller than the minimum wavelength of inhomogeneities that can arise during the reaction timescale via reaction-diffusion coupling.

In this well-mixed regime, reaction rates are homogenous across all parts of the container. The rate of change of the number of molecules $R_i$ with $i = 1, \ldots, n$ is in general a function of the numbers of molecules of all species and the container volume. This can be expressed as $\dot{R}_i = f_i(R_1, R_2, \ldots, R_n, V)$. Since all reactions occur homogeneously in space, the reaction rate per volume $g_i = f_i/V$ is the same everywhere. The system therefore has a scaling property

$$f_i(R_1\xi, R_2\xi, \ldots, R_n\xi, V\xi) = \xi f_i(R_1, R_2, \ldots, R_n, V) \quad . \tag{22}$$

Choosing $\xi = 1/V$ reveals that reaction rates per volume $g_i(r_1, r_2 ... r_n)$ are functions of the concentrations $r_i = R_i/V$ only. We thus find that the reaction fluxes $\dot{R}_i$ are extensive and obey

$$\dot{R}_i = V g_i(r_1, r_2 ... r_n) \quad . \tag{23}$$

Since the volume $V(R_1, \ldots, R_n)$ depends on composition and therefore varies, we find

$$\dot{r}_i = \frac{\dot{R}_i}{V} - \frac{R_i \dot{V}}{V^2} = g_i(r_1, r_2, \ldots, r_n) - r_i \frac{\dot{V}}{V} \quad . \tag{24}$$

Thus intensive condensate dynamics will arise whenever $\dot{V}/V$ is itself intensive and independent of volume. This is equivalent to

$$\dot{V} = V h(r_1, r_2, \ldots, r_n) \quad . \tag{25}$$

Before determining the volume growth rate $h$ we first note that $V$ is extensive according to

$$V(R_1\xi, R_2\xi, \ldots, R_n\xi) = \xi V(R_1, R_2, \ldots, R_n) \quad . \tag{26}$$

Taking a derivative with respect to $\xi$ and setting $\xi = 1$ we have

$$V(R_1, R_2, \ldots, R_n) = \sum_{i=1}^{n} R_i \frac{\partial V}{\partial R_i} \quad . \tag{27}$$

From equation 26 it follows that the effective molecular volumes

$$v_i(r_1, \ldots, r_n) = \frac{\partial V}{\partial R_i} \tag{28}$$

are intensive. Next, to determine the volume growth rate $h$ that leads to intensive chemical dynamics we have $h = \dot{V}/V$ which yields

$$h = \frac{1}{V} \sum_{i=1}^{n} \frac{\partial V}{\partial R_i} \frac{dR_i}{dt} + \frac{1}{V} \sum_{i=1}^{n} R_i \frac{d}{dt} (\frac{\partial V}{\partial R_i}) \quad , \tag{29}$$

which simplifies to

$$h = \sum_{i=1}^{n} g_i v_i + r_i \dot{v}_i \quad . \tag{30}$$

equation 27 ensures that the sum $\sum_{i=1}^{n} r_i \dot{v}_i = 0$ giving,

$$h = \sum_{i=1}^{n} g_i v_i \quad , \tag{31}$$

Finally, using equation 24, we also obtain a rate of concentration changes.

$$\dot{r_i} = g_i - r_i h \quad . \tag{32}$$

When $h$ is zero, that is for fixed volume systems, we recover conventional mass action kinetics of concentrations $\dot{r_i} = g_i$.

Note that the volume growth rate $h$ and the concentration dynamics $\dot{r_i}$ are intensive if the system is well mixed with extensive total volume, and therefore both the volume fluxes and reaction fluxes are extensive.

Thus we learn, that while fixed volume containers need only be well-mixed in order to generate intensive chemical dynamics, a variable-volume container additionally requires that the rate of volume change be determined by an intensive per-volume generation rate ($h$). For cortical condensates this is true, because internal molecular amounts change via homogenous mass action kinetics and simultaneously determine the volume.

### 9 Chemical reactions generate an effective critical condensate size

In both control and RNAi datasets, the nullclines for WSP-1 and F-actin are linear and pass through the origin (Figure 2 C, 3 C). The WSP-1 nullcline may be interpreted as a set of critical sizes for WSP-1 growth - that is, to note that for WSP-1 amounts above this line, assemblies add new WSP-1 molecules while below this line they lose WSP-1 molecules just as classical nucleation theory exhibits a nucleation barrier peaked at a critical size. The linear nullcline then implies that the critical size for WSP-1 growth increases linearly with internal F-actin content rather than being fixed by WSP-1 - WSP-1 interaction strength and external concentrations as it would be for the nucleation and growth of single component WSP-1 droplets from their supersaturated vapor. Additionally, the observation that the nullcline passes through the origin means that in the absence of F-actin, an all WSP-1 assembly exhibits a zero critical size. Together with the fact that each WSP-1 molecule contributes the same self-recruitment rate independent of the other WSP-1 molecules in the condensate (self-recruitment rates $\sim k_r W$), these observations suggest that the critical size for WSP-1 growth is generated by reaction kinetics rather than by the thermodynamics of nucleation and growth.

**Supplementary Movie caption**

**Movie S1** *In utero* confocal spinning disc imaging of *C. elegans* oocyte to embryo transition. Lifeact::mKate labeling F-actin in magenta, Histones H2B::GFP labeled in green. Scale bar, 10 $\mu$m. Time stamp (min:sec).

**Movie S2** Formation of an actomyosin cortex in an isolated *C. elegans* oocyte. Lifeact::mKate labeling F-actin in magenta, Non-muscle myosin NMY-2::GFP labeled in green. Scale bar, 10 $\mu$m. Time stamp (min:sec).

**Movie S3** Cortical assemblies in a *C. elegans* oocyte undergoing actomyosin cortex formation. Lifeact::mKate labeling F-actin in magenta, endogenously labeled WSP-1::GFP in green. Scale bar, 10 $\mu$m. Time stamp (min:sec).

**Movie S4** Cortical assemblies in a *C.elegans* oocyte contain WSP-1 and ARX-2. Endogenously labeled WSP-1::GFP in green, and endogenously labeled ARX-2::mCherry in blue. Scale bar, 10 $\mu$m. Time stamp (min:sec).

**Movie S5** Oocyte severely depleted of ARX-2 after 20 hours of RNAi feeding. Lifeact::mKate labeling F-actin in magenta, Non-muscle myosin NMY-2::GFP labeled in green. Scale bar, 10 $\mu$m. Time stamp (min:sec).

**Movie S6** Oocyte mildly depleted of ARX-2 after 19 hours of RNAi feeding. Lifeact::mKate labeling F-actin in magenta, endogenous WSP-1::GFP labeled in green. Scale bar, 10 $\mu$m. Time stamp (min:sec).

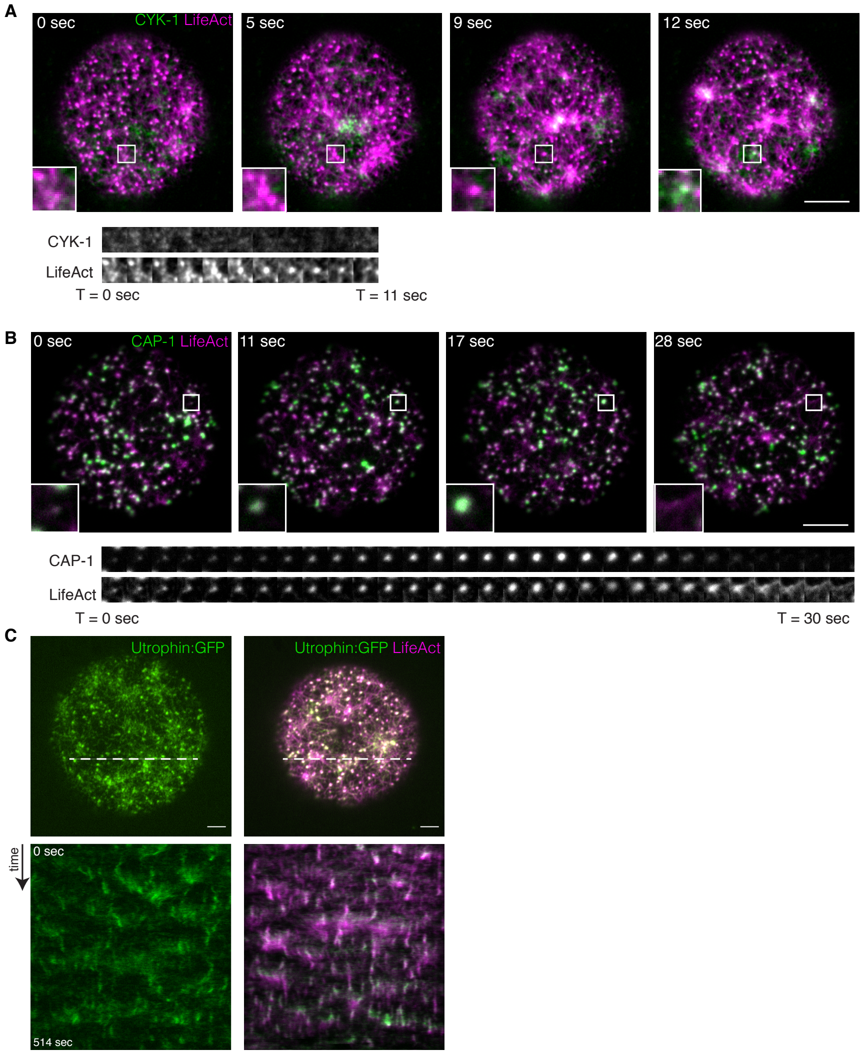


**Supplementary figure S1 : F-actin puncta form independently of Lifeact labeling, CAP-1 but not Formin CYK-1 is enriched in the F-actin puncta.**

**Figure S1 Caption: A**. CYK-1/Formin does not colocalize with Lifeact condensates, but is found in contractile actomyosin pulses just prior to contractions. **B**. Capping protein CAP-1 localizes to F-actin condensates. **C**. Alternative F-actin labeling (Utrophin) does not affect condensate dynamics. Scale bar is 10 $\mu$ m.

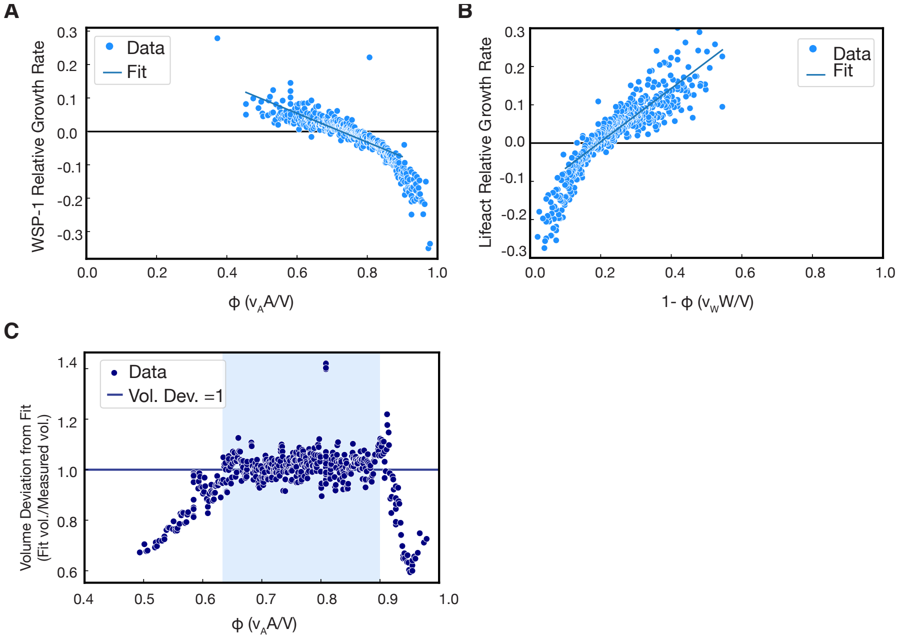


**Supplementary figure S2 : Volume dependence on molecular content and the linear dependence of relative growth rate on stoichiometry**

**Figure S2 Caption: A, B**. Linear dependence of WSP-1 (**A**)and F-actin (**B**) relative growth rate on stoichiometry for control oocytes. **C**. Deviation of measured volume from linear combination of WSP-1 and F-actin intensities for F-actin volume fraction $\phi <\sim .6$ and $\phi >\sim .9$. Note the deviation from linearity in **A, B**. coincides with the range of volume fractions where measured volume deviates from $V = v_A A + v_W W$. As shown in Figure 3 E,F of main text and in Figure S5 below, data from the ARX-2 RNAi oocytes occupies an experimental stoichiometry range with no deviation. Also see supplementary section 2.

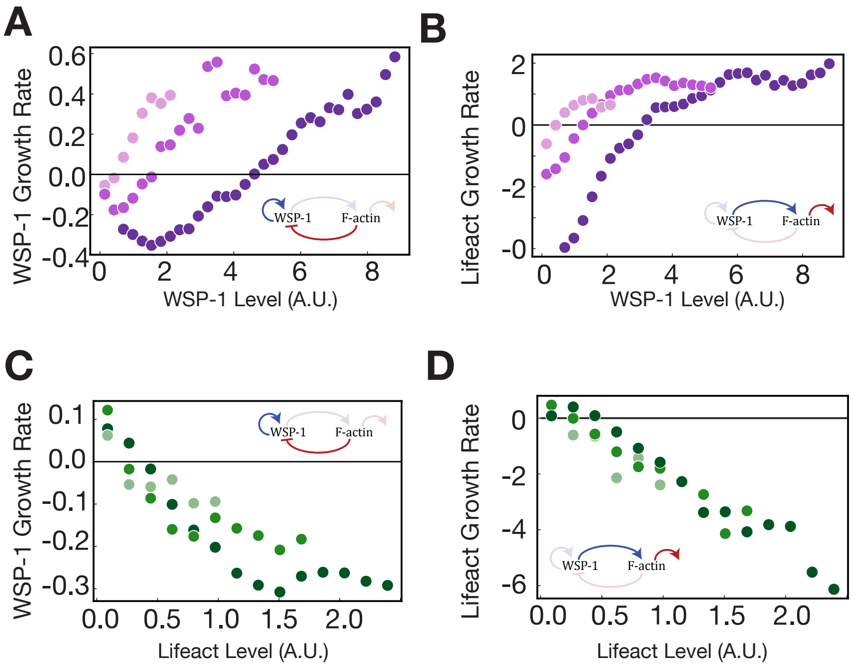


**Supplementary figure S3 : Mass flux phase portrait slices corroborate the empirical growth laws.**

**Figure S3 Caption: A,B**. WSP-1 **(A)** and F-actin **(B)** growth rates as a function of WSP-1 at constant levels of Lifeact (lowest three lifeact bins of Figure 2 C). **C, D**. WSP-1 **(C)** and F-actin **(D)** growth rates as a function of F-actin at constant levels of WSP-1 (Lowest three WSP-1 bins of Figure 2 C). Insets in **A-D** indicate feedback processes from Figure 2E that contribute in each case.

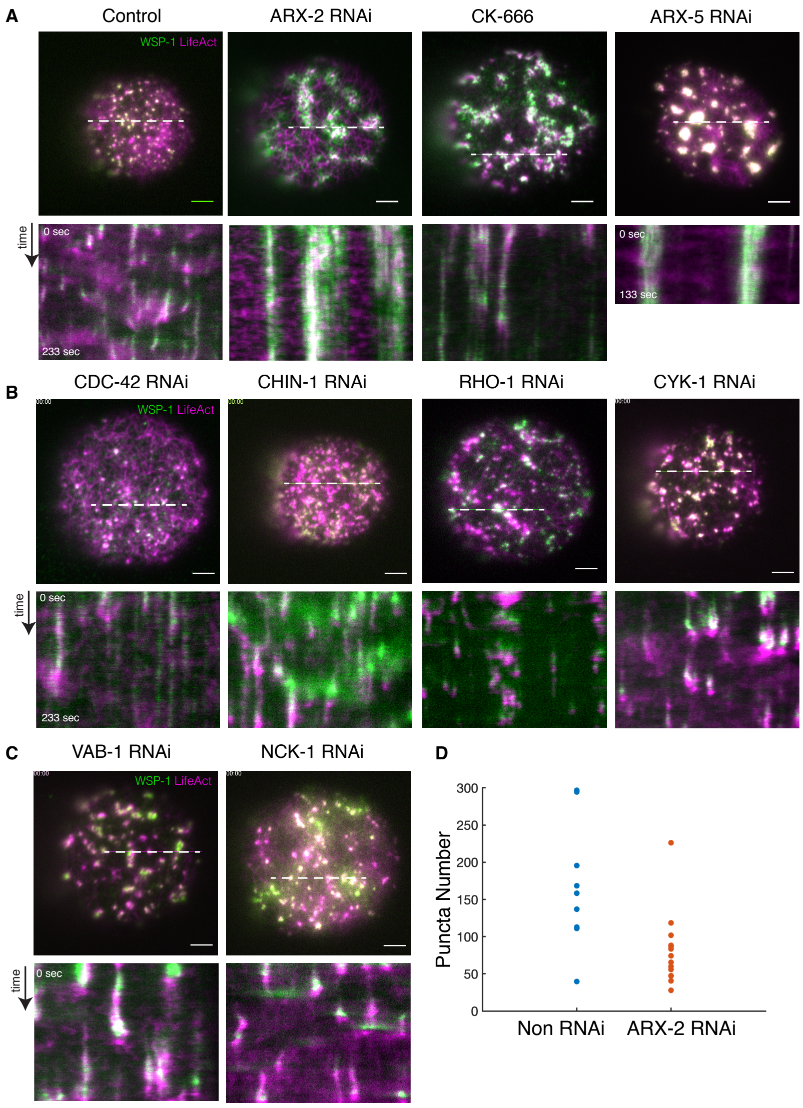


**Supplementary figure S4 : Genetic perturbations of WSP-1 and Formin based nucleation pathways**

**Figure S4 Caption:** **A**. Representative control oocyte (n=15), oocyte from C.elegans after 20 hours ARX-2 feeding (n=9), oocyte treated with 10 $\mu$M CK-666 (n = 4) to specifically inhibit Arp2/3 branching reaction and oocytes depleted of ARX-5 (ArpC3) (n = 4). Kymographs draw from dotted lines. **B**. RNAi depletion of CDC-42 and CHIN-1 resulted in altered levels of WSP-1 at the cortex in oocytes while maintaing dynamic condensates. Knockdown of RHO-1 and CYK-1 resulted in no observable changes in condensate dynamics. **C**. Knockdown of NCK-1 or the NCK-1 activator VAB-1 resulted in no observable changes in condensate dynamics. Scale bar = 5 $\mu$ m. **D**. ARX-2 depleted oocytes have reduced number of condensates compared to control oocytes.

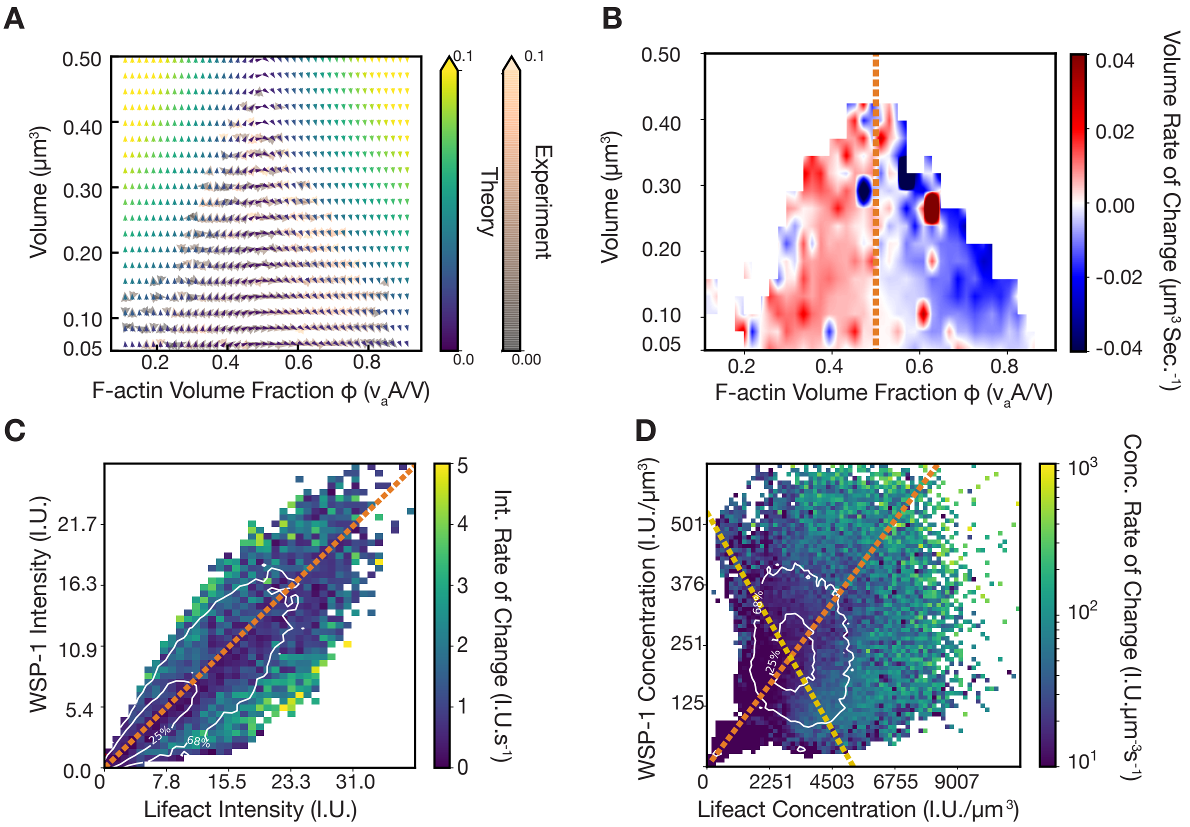


**Supplementary figure S5 : Kinetic slowdown (following Figure 4) in oocytes subjected to ARX-2 RNAi**

**Figure S5 caption:** **A**. Mass flux phase portrait in volume-volume fraction coordinates for both data and growth laws showing the close match to ARX-2 RNAi data in a second coordinate system, as in main text Figure 4. **B**. Heat map representing the rate of change of volume in ARX-2 RNAi conditions as a function of instantaneous volume and F-actin volume fraction. The F-actin volume fraction ∼0.5 (orange line in **B** and corresponding to the orange line at intensity stoichiometry ∼ 0.59 in **C, D**) coincides with a switch from growing volumes (assembly) to shrinking volumes (disassembly) **C**. Heat map representing the magnitude of change of F-actin and WSP-1 amounts per unit time as a function of instantaneous WSP-1 and F-actin amounts in ARX-2 RNAi conditions. Orange line depicts the stoichiometry ∼0.59 preferentially maintained by the ensemble (main text Figure 3D) and coincides with the slowest rates of change. **D**. Heat map representing the rate of change of F-actin and WSP-1 concentrations as a function of instantaneous concentrations in ARX-2 RNAi conditions. The contours depict the most commonly occupied concentration values and reflect the preferential maintenance of a pair of concentrations. This concentration pair lies on both the line of contant total density (determined by the volume constraint $V = v_A A + v_W W$) - yellow line, and the stoichiometry of slowest concentration changes - orange line.

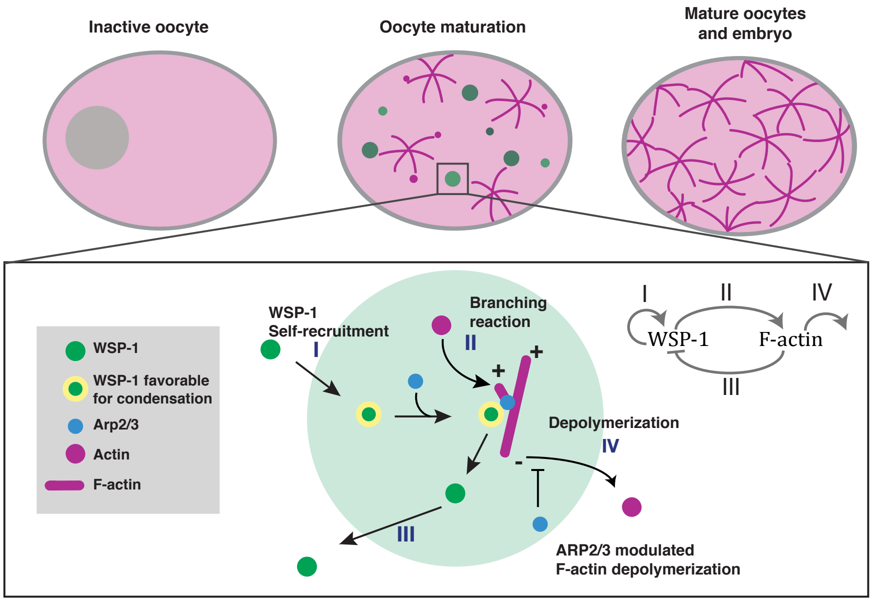


**Supplementary figure S6 : Biochemical interpretations of kinetic coefficients as suggested by their change on ARX-2 RNAi**

**Figure S6 caption:** Schematic of oocyte cortex formation through an intermediate stage with WSP-1, ARP2/3, and F-actin condensates. The four processes that underly the dynamic instability of condensates are WSP-1 self-recruitment, branching-dependent WSP-1 loss, branching-dependent F-actin growth, and F-actin loss (depolymerization or severing modulated by ARP2/3). Analysis of the effect of ARX-2 RNAi on the kinetic coefficients yields consistency with established biochemical processes as indicated.
